## Supplemental Figures for "NPRL2 gene therapy induces effective antitumor immunity in KRAS/STK11 mutant anti-PD1 resistant metastatic non-small cell lung cancer (NSCLC) in a humanized mouse model"

### Slide 1
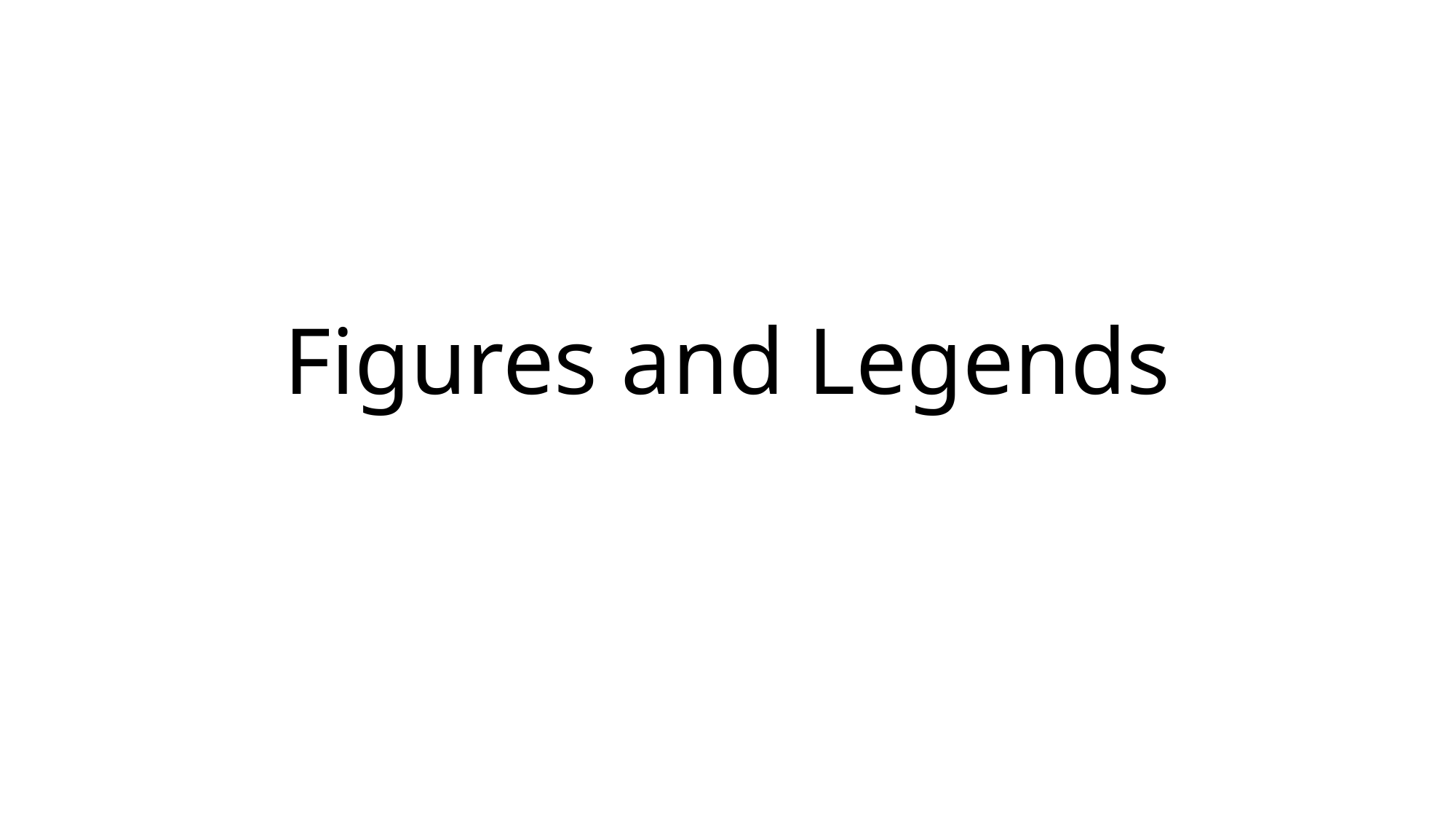

#### Figures and Legends

### Slide 2
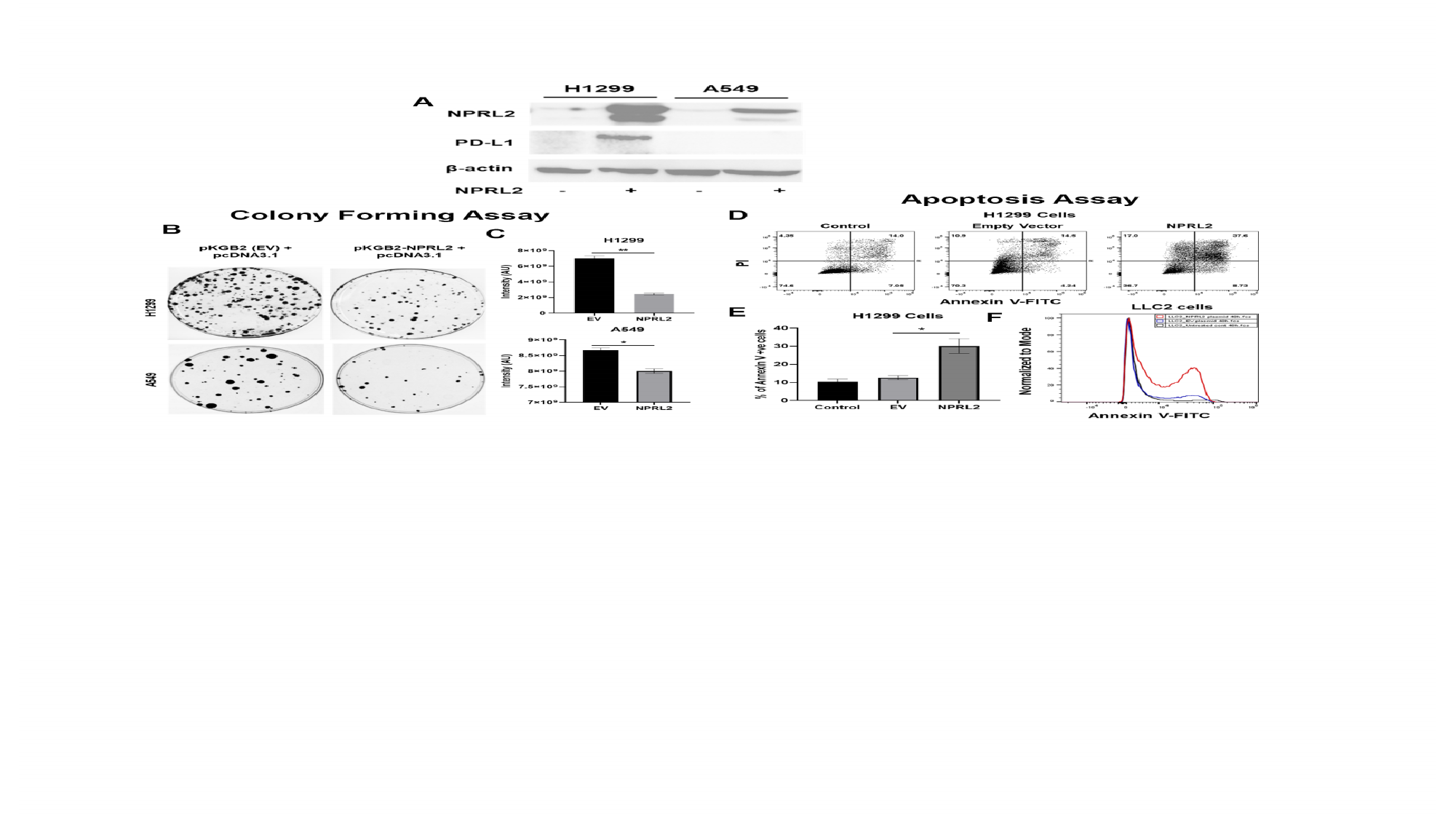

### Slide 3
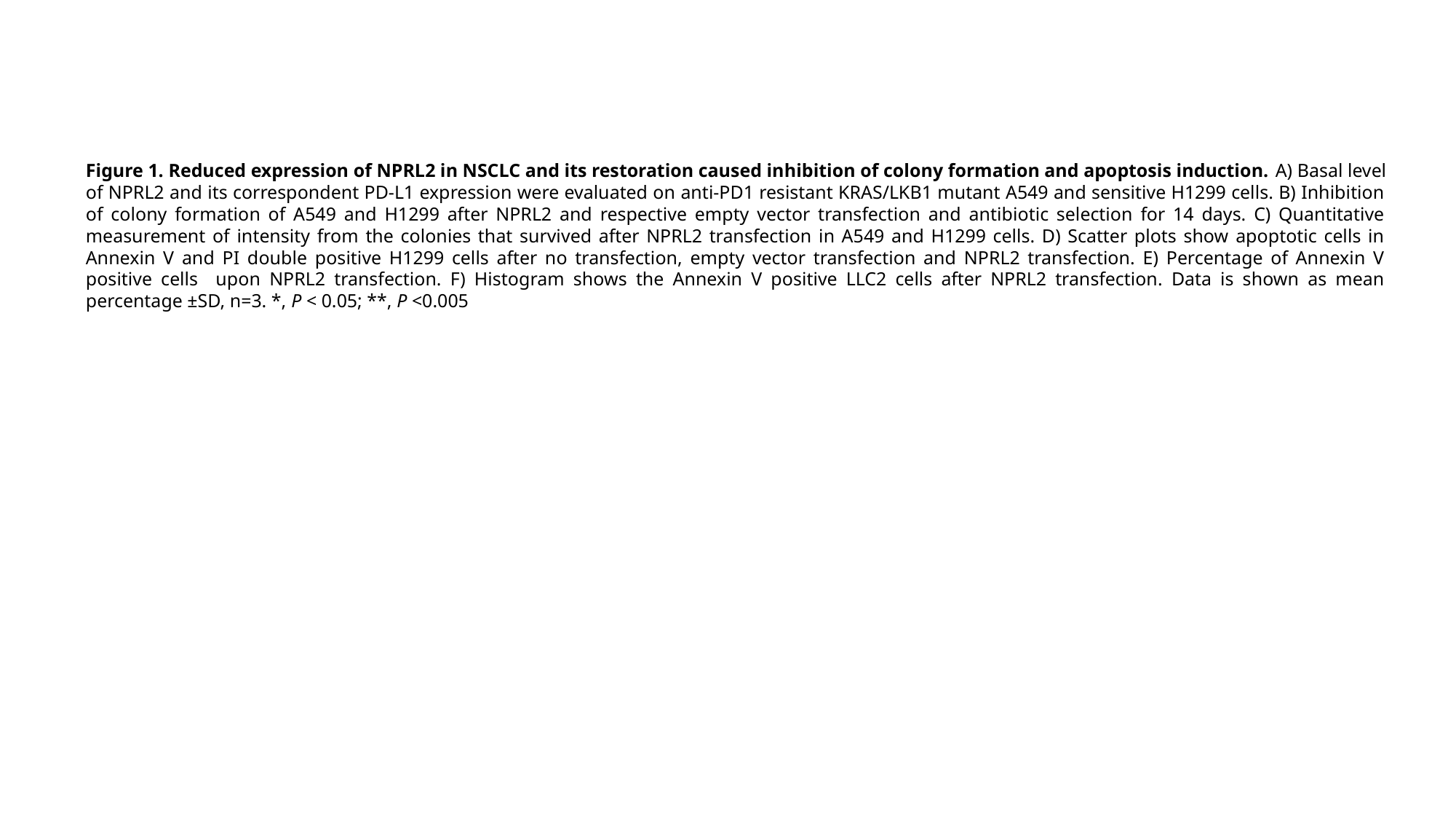

Figure 1. Reduced expression of NPRL2 in NSCLC and its restoration caused inhibition of colony formation and apoptosis induction. A) Basal level of NPRL2 and its correspondent PD-L1 expression were evaluated on anti-PD1 resistant KRAS/LKB1 mutant A549 and sensitive H1299 cells. B) Inhibition of colony formation of A549 and H1299 after NPRL2 and respective empty vector transfection and antibiotic selection for 14 days. C) Quantitative measurement of intensity from the colonies that survived after NPRL2 transfection in A549 and H1299 cells. D) Scatter plots show apoptotic cells in Annexin V and PI double positive H1299 cells after no transfection, empty vector transfection and NPRL2 transfection. E) Percentage of Annexin V positive cells upon NPRL2 transfection. F) Histogram shows the Annexin V positive LLC2 cells after NPRL2 transfection. Data is shown as mean percentage ±SD, n=3. *, P < 0.05; **, P <0.005

### Slide 4
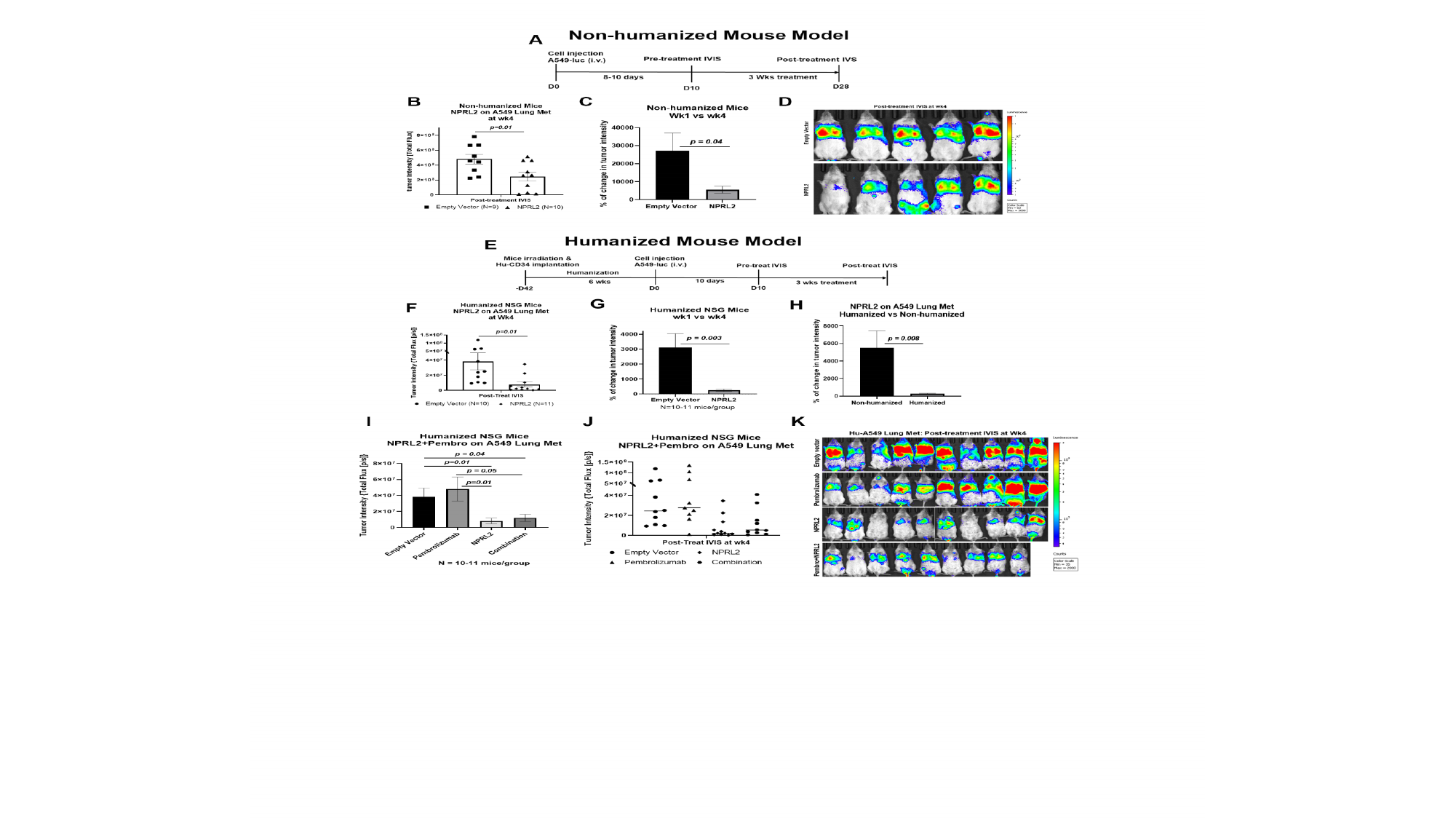

### Slide 5
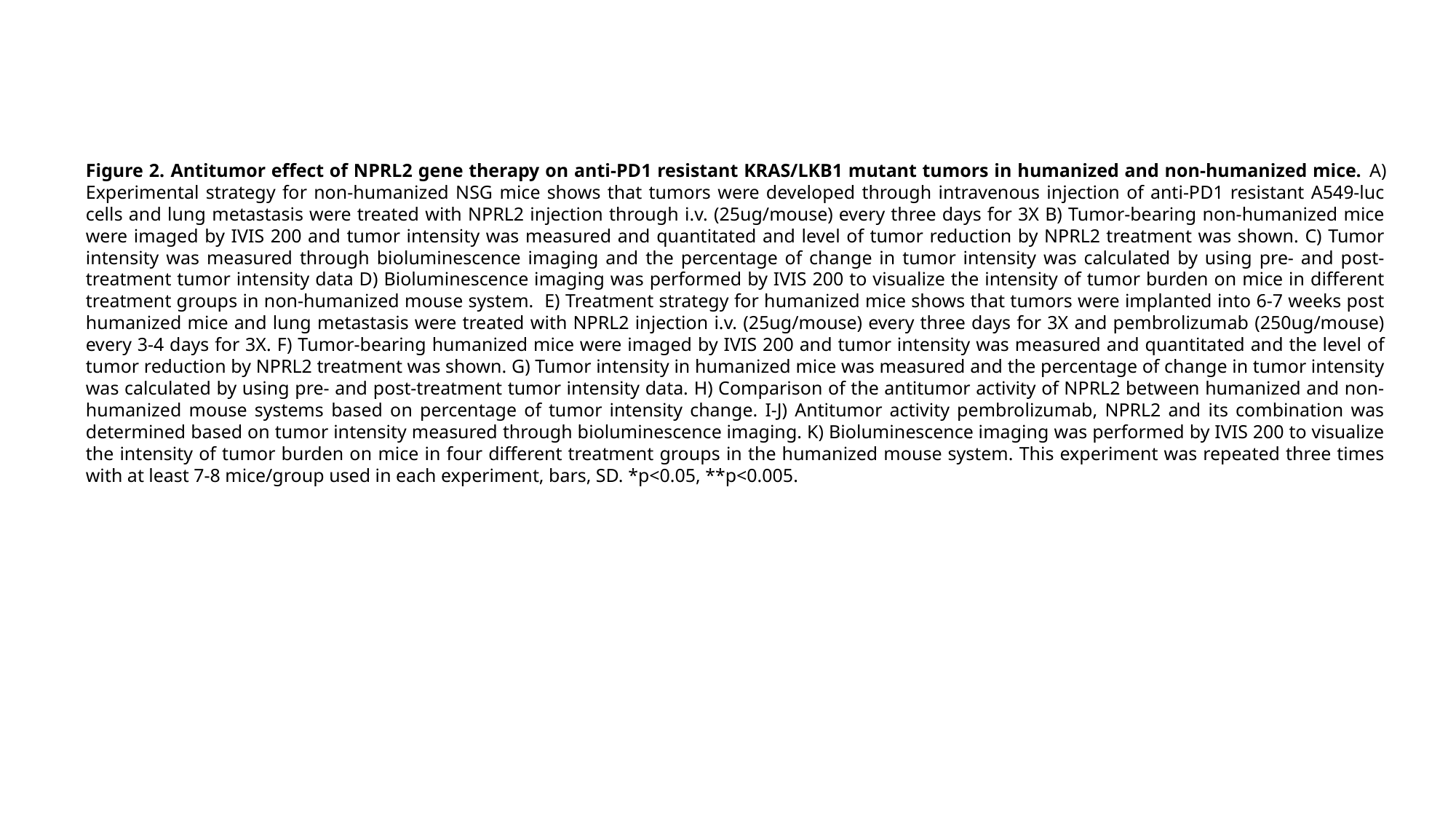

Figure 2. Antitumor effect of NPRL2 gene therapy on anti-PD1 resistant KRAS/LKB1 mutant tumors in humanized and non-humanized mice. A) Experimental strategy for non-humanized NSG mice shows that tumors were developed through intravenous injection of anti-PD1 resistant A549-luc cells and lung metastasis were treated with NPRL2 injection through i.v. (25ug/mouse) every three days for 3X B) Tumor-bearing non-humanized mice were imaged by IVIS 200 and tumor intensity was measured and quantitated and level of tumor reduction by NPRL2 treatment was shown. C) Tumor intensity was measured through bioluminescence imaging and the percentage of change in tumor intensity was calculated by using pre- and post-treatment tumor intensity data D) Bioluminescence imaging was performed by IVIS 200 to visualize the intensity of tumor burden on mice in different treatment groups in non-humanized mouse system. E) Treatment strategy for humanized mice shows that tumors were implanted into 6-7 weeks post humanized mice and lung metastasis were treated with NPRL2 injection i.v. (25ug/mouse) every three days for 3X and pembrolizumab (250ug/mouse) every 3-4 days for 3X. F) Tumor-bearing humanized mice were imaged by IVIS 200 and tumor intensity was measured and quantitated and the level of tumor reduction by NPRL2 treatment was shown. G) Tumor intensity in humanized mice was measured and the percentage of change in tumor intensity was calculated by using pre- and post-treatment tumor intensity data. H) Comparison of the antitumor activity of NPRL2 between humanized and non-humanized mouse systems based on percentage of tumor intensity change. I-J) Antitumor activity pembrolizumab, NPRL2 and its combination was determined based on tumor intensity measured through bioluminescence imaging. K) Bioluminescence imaging was performed by IVIS 200 to visualize the intensity of tumor burden on mice in four different treatment groups in the humanized mouse system. This experiment was repeated three times with at least 7-8 mice/group used in each experiment, bars, SD. *p<0.05, **p<0.005.

### Slide 6
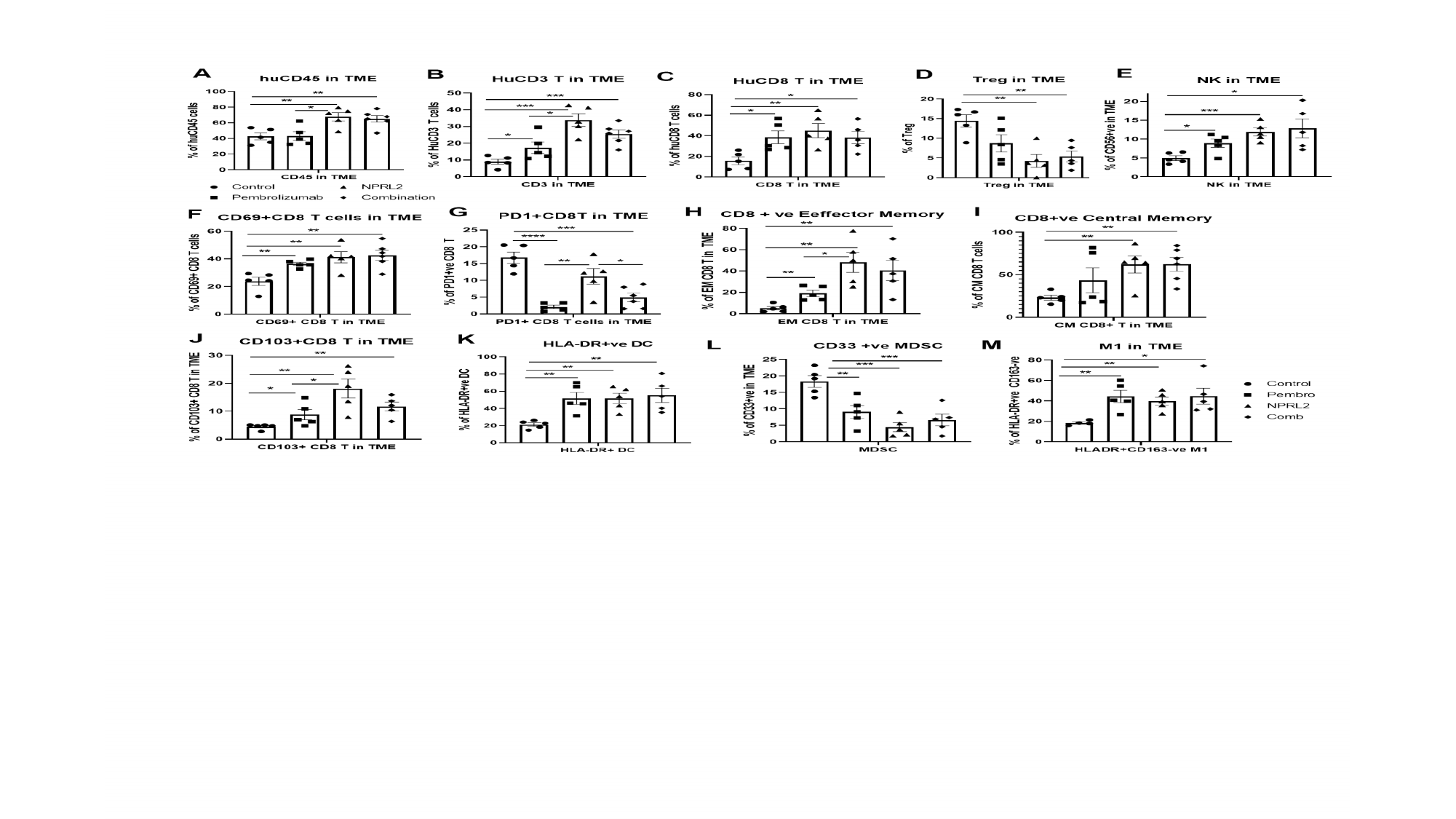

### Slide 7
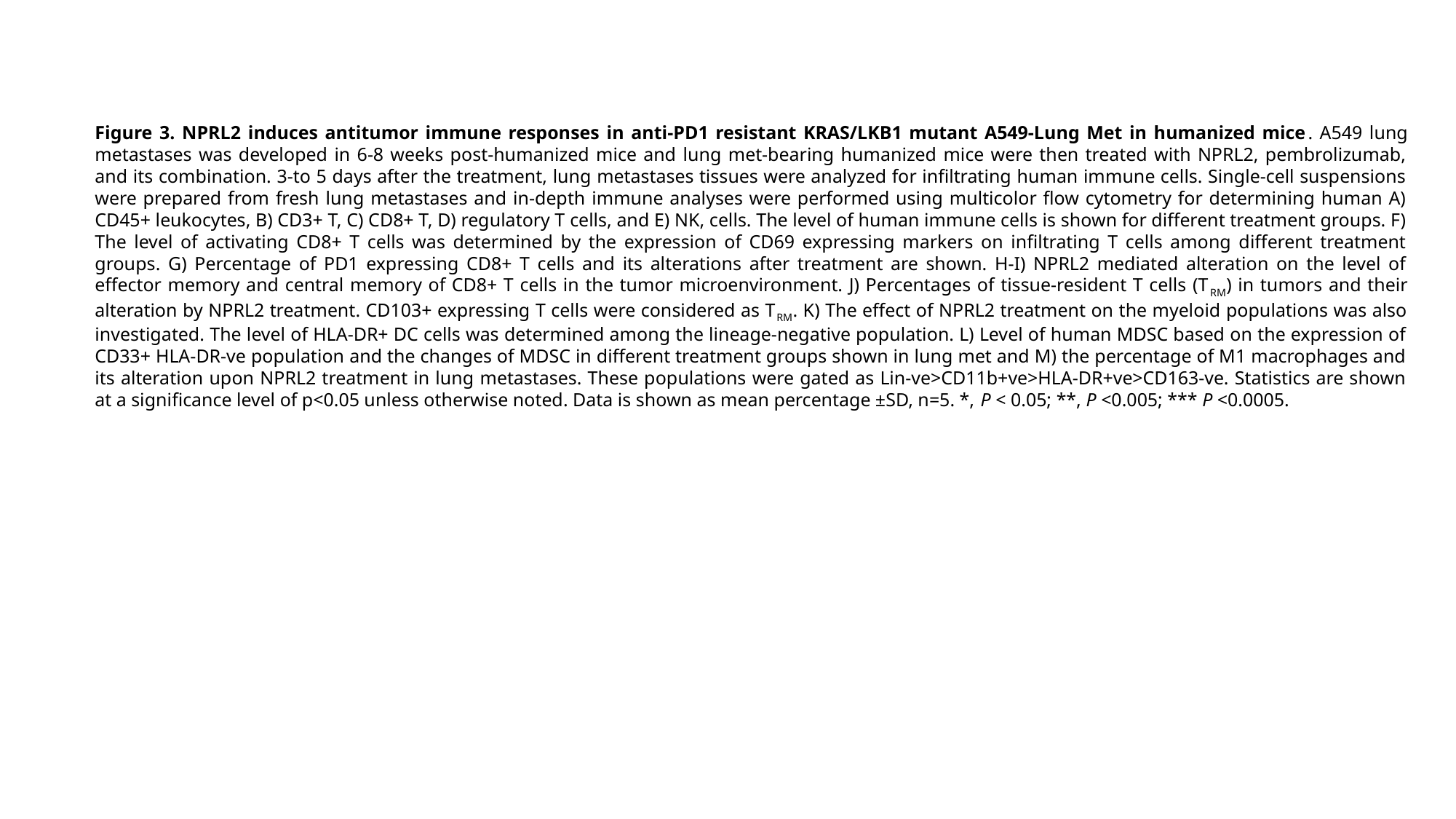

Figure 3. NPRL2 induces antitumor immune responses in anti-PD1 resistant KRAS/LKB1 mutant A549-Lung Met in humanized mice. A549 lung metastases was developed in 6-8 weeks post-humanized mice and lung met-bearing humanized mice were then treated with NPRL2, pembrolizumab, and its combination. 3-to 5 days after the treatment, lung metastases tissues were analyzed for infiltrating human immune cells. Single-cell suspensions were prepared from fresh lung metastases and in-depth immune analyses were performed using multicolor flow cytometry for determining human A) CD45+ leukocytes, B) CD3+ T, C) CD8+ T, D) regulatory T cells, and E) NK, cells. The level of human immune cells is shown for different treatment groups. F) The level of activating CD8+ T cells was determined by the expression of CD69 expressing markers on infiltrating T cells among different treatment groups. G) Percentage of PD1 expressing CD8+ T cells and its alterations after treatment are shown. H-I) NPRL2 mediated alteration on the level of effector memory and central memory of CD8+ T cells in the tumor microenvironment. J) Percentages of tissue-resident T cells (T_RM_) in tumors and their alteration by NPRL2 treatment. CD103+ expressing T cells were considered as T_RM_. K) The effect of NPRL2 treatment on the myeloid populations was also investigated. The level of HLA-DR+ DC cells was determined among the lineage-negative population. L) Level of human MDSC based on the expression of CD33+ HLA-DR-ve population and the changes of MDSC in different treatment groups shown in lung met and M) the percentage of M1 macrophages and its alteration upon NPRL2 treatment in lung metastases. These populations were gated as Lin-ve>CD11b+ve>HLA-DR+ve>CD163-ve. Statistics are shown at a significance level of p<0.05 unless otherwise noted. Data is shown as mean percentage ±SD, n=5. *, P < 0.05; **, P <0.005; *** P <0.0005.

### Slide 8
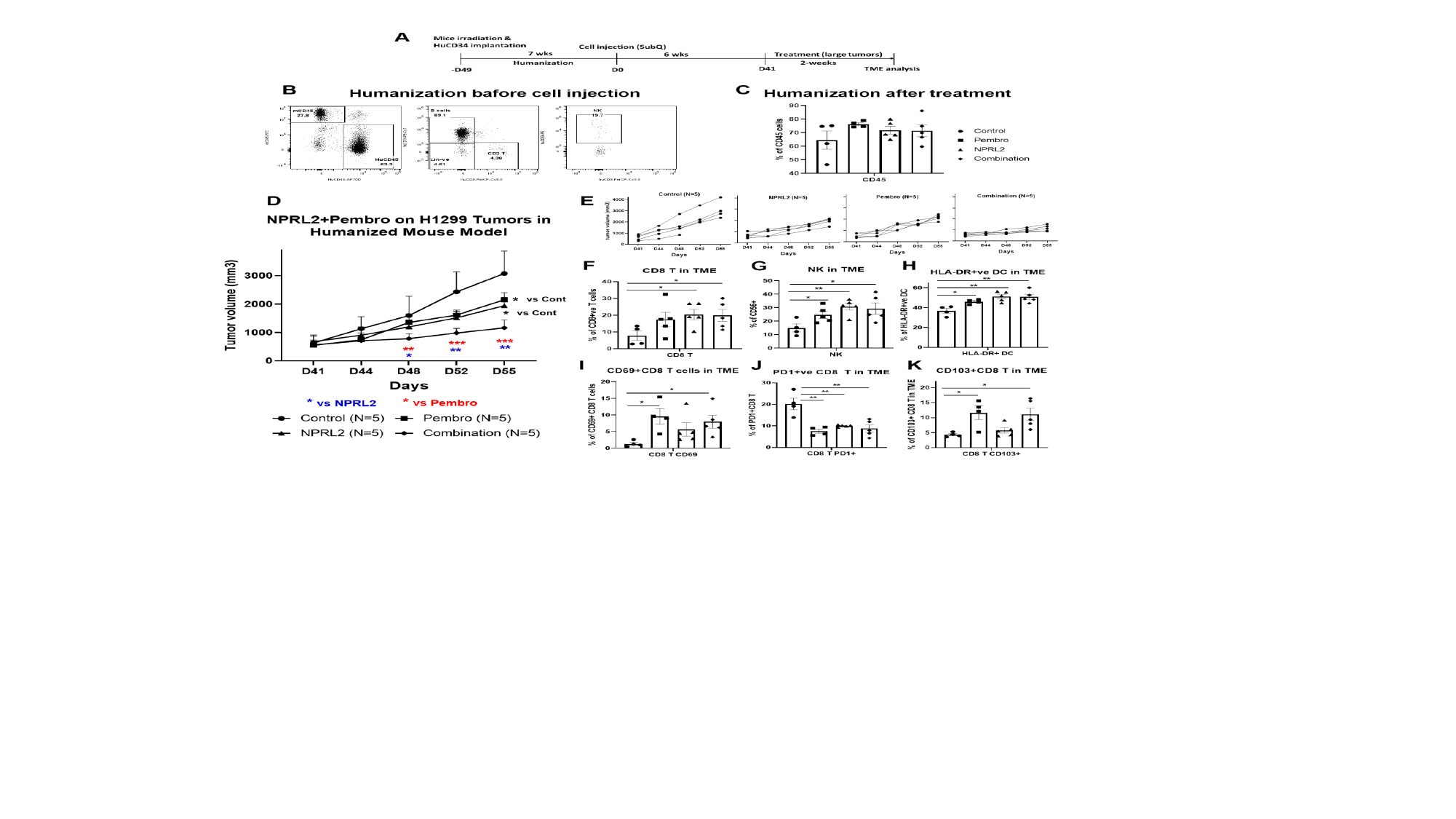

### Slide 9
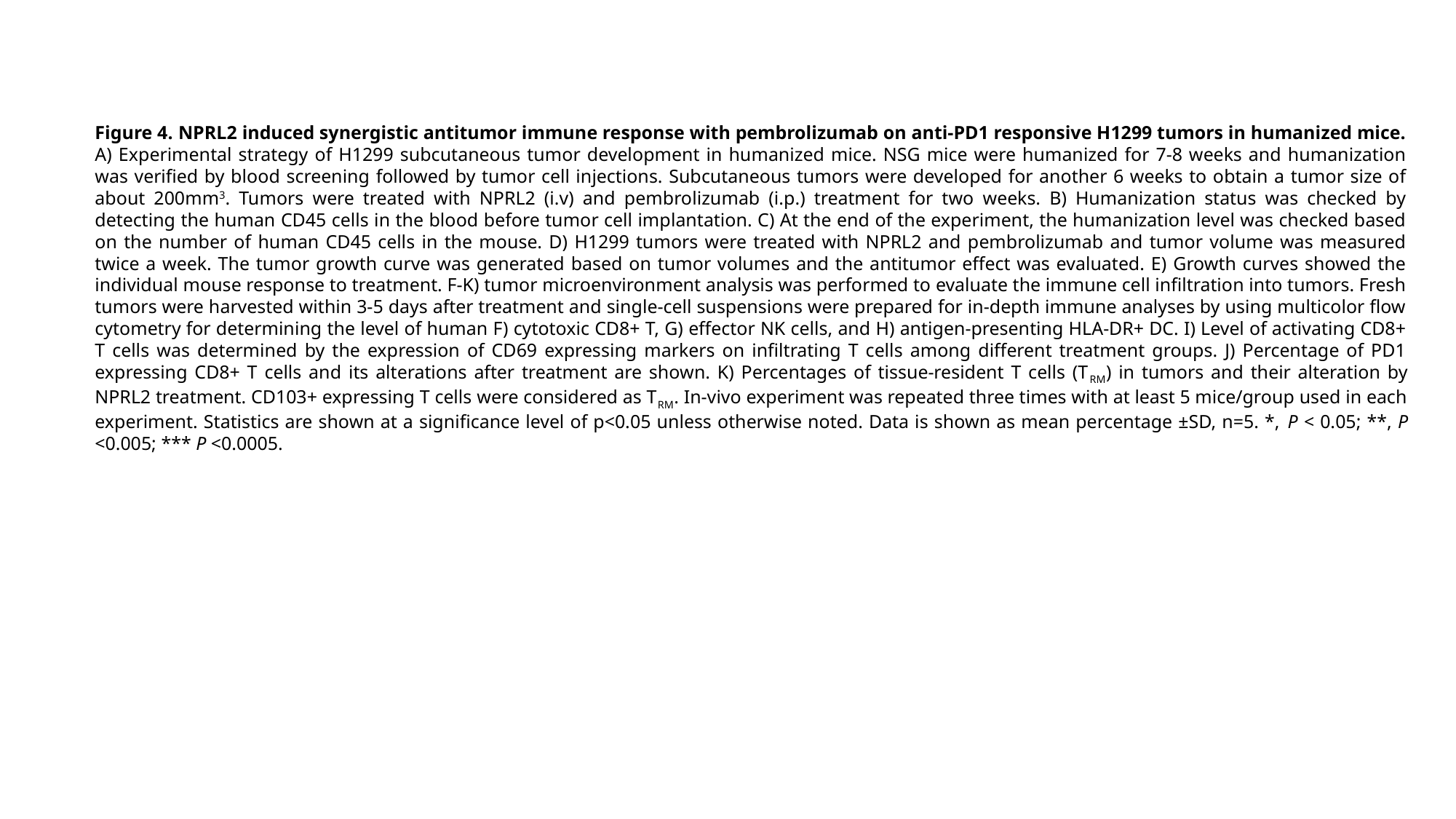

Figure 4. NPRL2 induced synergistic antitumor immune response with pembrolizumab on anti-PD1 responsive H1299 tumors in humanized mice. A) Experimental strategy of H1299 subcutaneous tumor development in humanized mice. NSG mice were humanized for 7-8 weeks and humanization was verified by blood screening followed by tumor cell injections. Subcutaneous tumors were developed for another 6 weeks to obtain a tumor size of about 200mm^3^. Tumors were treated with NPRL2 (i.v) and pembrolizumab (i.p.) treatment for two weeks. B) Humanization status was checked by detecting the human CD45 cells in the blood before tumor cell implantation. C) At the end of the experiment, the humanization level was checked based on the number of human CD45 cells in the mouse. D) H1299 tumors were treated with NPRL2 and pembrolizumab and tumor volume was measured twice a week. The tumor growth curve was generated based on tumor volumes and the antitumor effect was evaluated. E) Growth curves showed the individual mouse response to treatment. F-K) tumor microenvironment analysis was performed to evaluate the immune cell infiltration into tumors. Fresh tumors were harvested within 3-5 days after treatment and single-cell suspensions were prepared for in-depth immune analyses by using multicolor flow cytometry for determining the level of human F) cytotoxic CD8+ T, G) effector NK cells, and H) antigen-presenting HLA-DR+ DC. I) Level of activating CD8+ T cells was determined by the expression of CD69 expressing markers on infiltrating T cells among different treatment groups. J) Percentage of PD1 expressing CD8+ T cells and its alterations after treatment are shown. K) Percentages of tissue-resident T cells (T_RM_) in tumors and their alteration by NPRL2 treatment. CD103+ expressing T cells were considered as T_RM_. In-vivo experiment was repeated three times with at least 5 mice/group used in each experiment. Statistics are shown at a significance level of p<0.05 unless otherwise noted. Data is shown as mean percentage ±SD, n=5. *, P < 0.05; **, P <0.005; *** P <0.0005.

### Slide 10
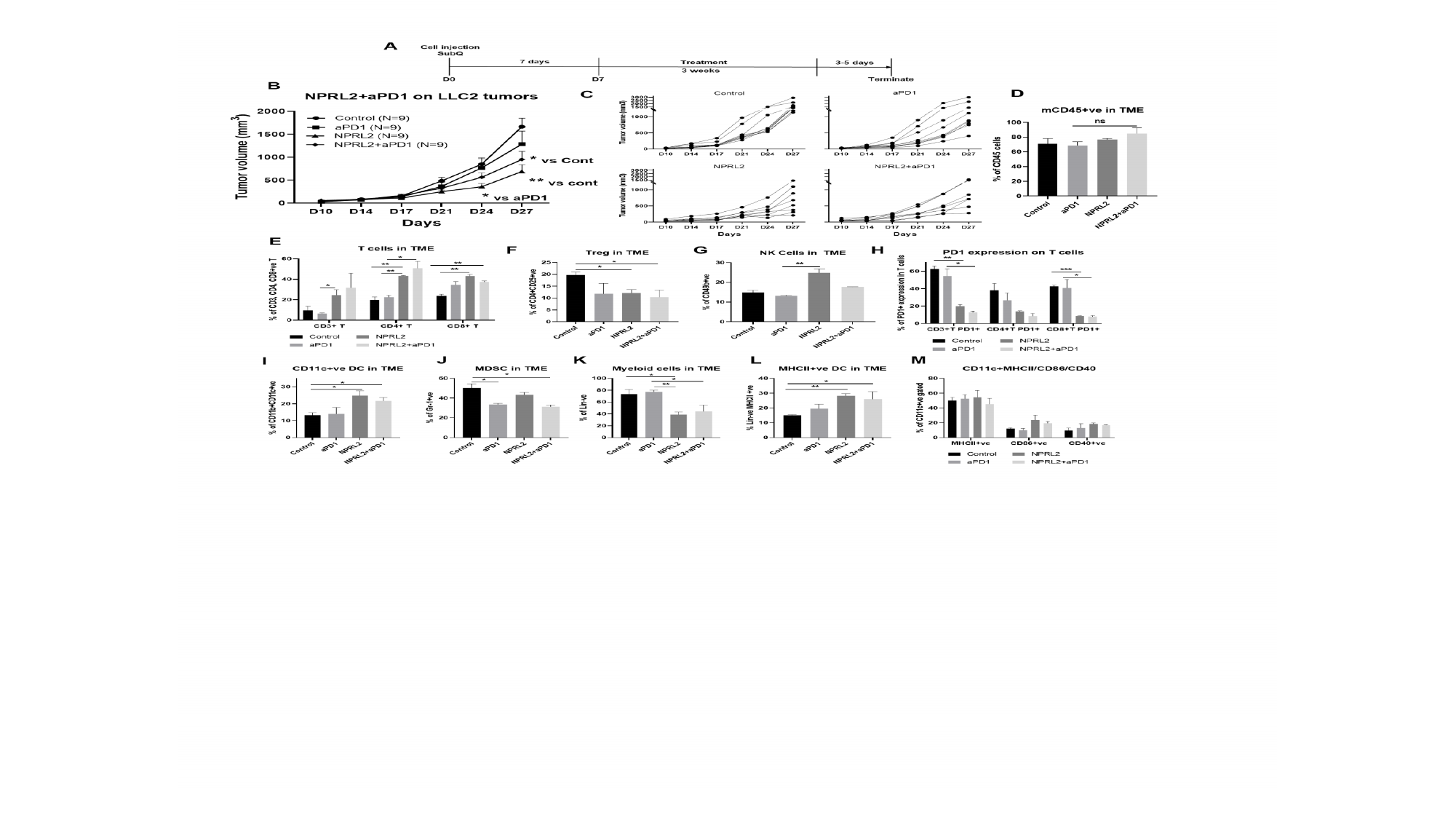

### Slide 11
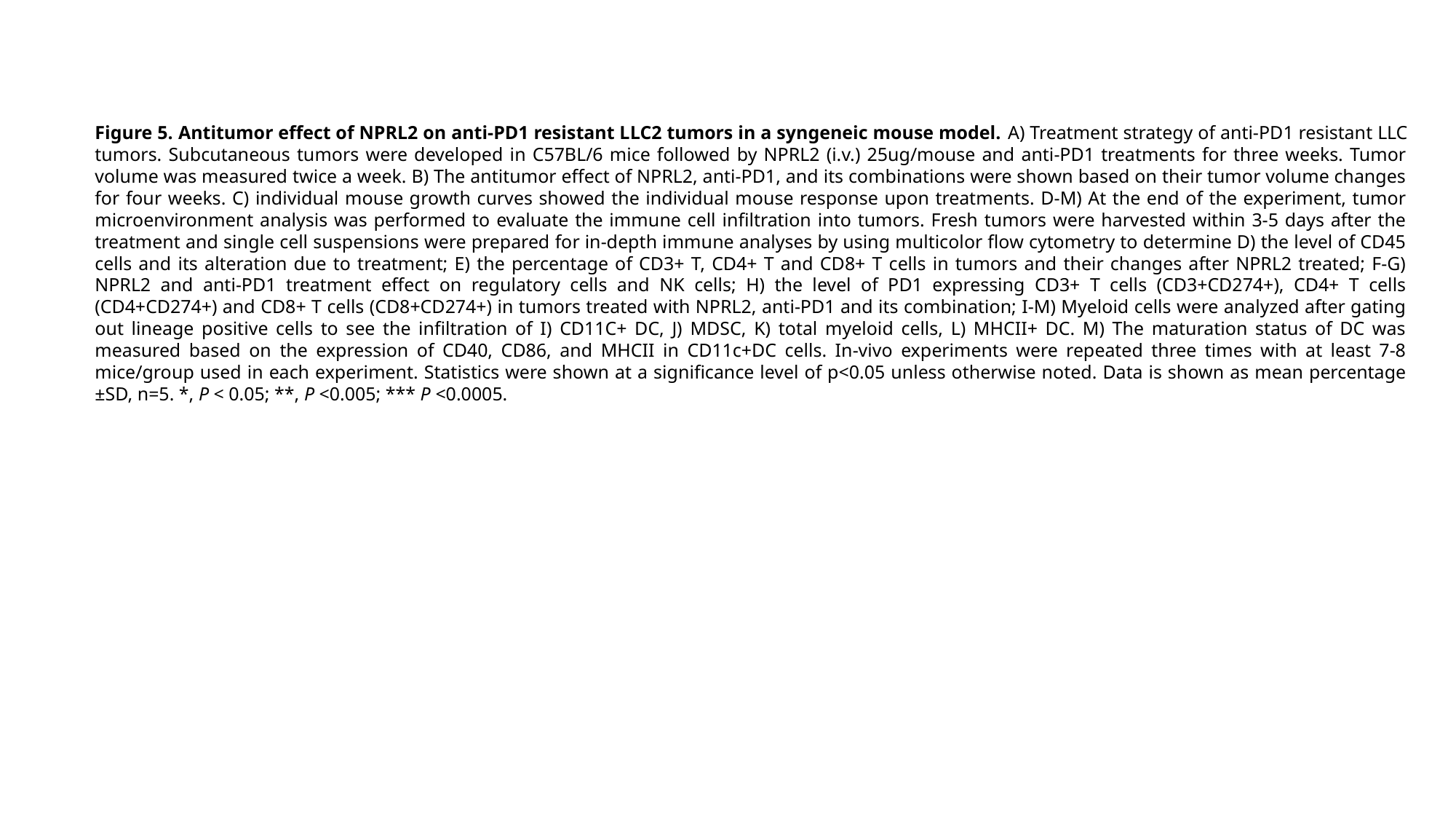

Figure 5. Antitumor effect of NPRL2 on anti-PD1 resistant LLC2 tumors in a syngeneic mouse model. A) Treatment strategy of anti-PD1 resistant LLC tumors. Subcutaneous tumors were developed in C57BL/6 mice followed by NPRL2 (i.v.) 25ug/mouse and anti-PD1 treatments for three weeks. Tumor volume was measured twice a week. B) The antitumor effect of NPRL2, anti-PD1, and its combinations were shown based on their tumor volume changes for four weeks. C) individual mouse growth curves showed the individual mouse response upon treatments. D-M) At the end of the experiment, tumor microenvironment analysis was performed to evaluate the immune cell infiltration into tumors. Fresh tumors were harvested within 3-5 days after the treatment and single cell suspensions were prepared for in-depth immune analyses by using multicolor flow cytometry to determine D) the level of CD45 cells and its alteration due to treatment; E) the percentage of CD3+ T, CD4+ T and CD8+ T cells in tumors and their changes after NPRL2 treated; F-G) NPRL2 and anti-PD1 treatment effect on regulatory cells and NK cells; H) the level of PD1 expressing CD3+ T cells (CD3+CD274+), CD4+ T cells (CD4+CD274+) and CD8+ T cells (CD8+CD274+) in tumors treated with NPRL2, anti-PD1 and its combination; I-M) Myeloid cells were analyzed after gating out lineage positive cells to see the infiltration of I) CD11C+ DC, J) MDSC, K) total myeloid cells, L) MHCII+ DC. M) The maturation status of DC was measured based on the expression of CD40, CD86, and MHCII in CD11c+DC cells. In-vivo experiments were repeated three times with at least 7-8 mice/group used in each experiment. Statistics were shown at a significance level of p<0.05 unless otherwise noted. Data is shown as mean percentage ±SD, n=5. *, P < 0.05; **, P <0.005; *** P <0.0005.

### Slide 12
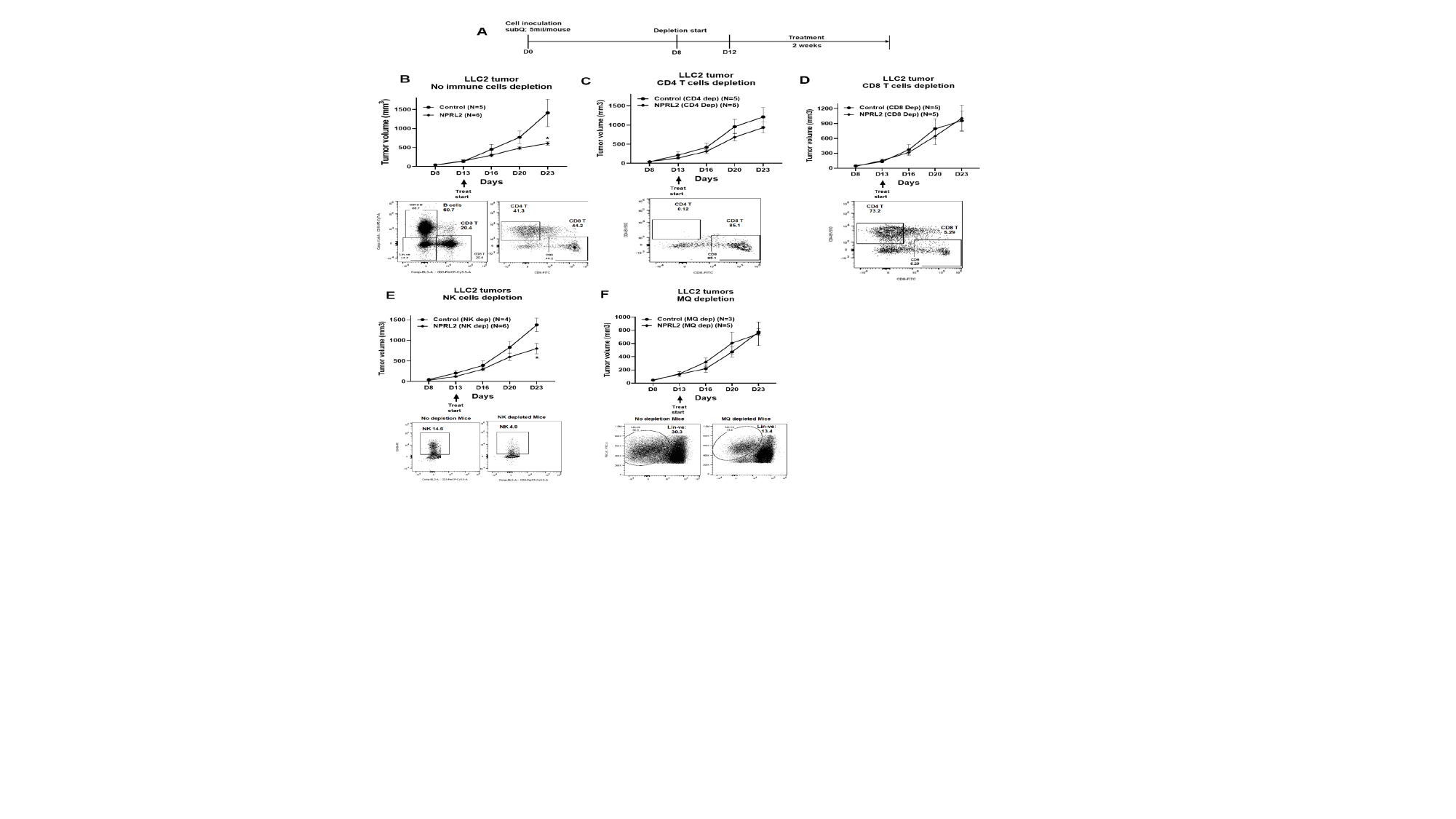

### Slide 13
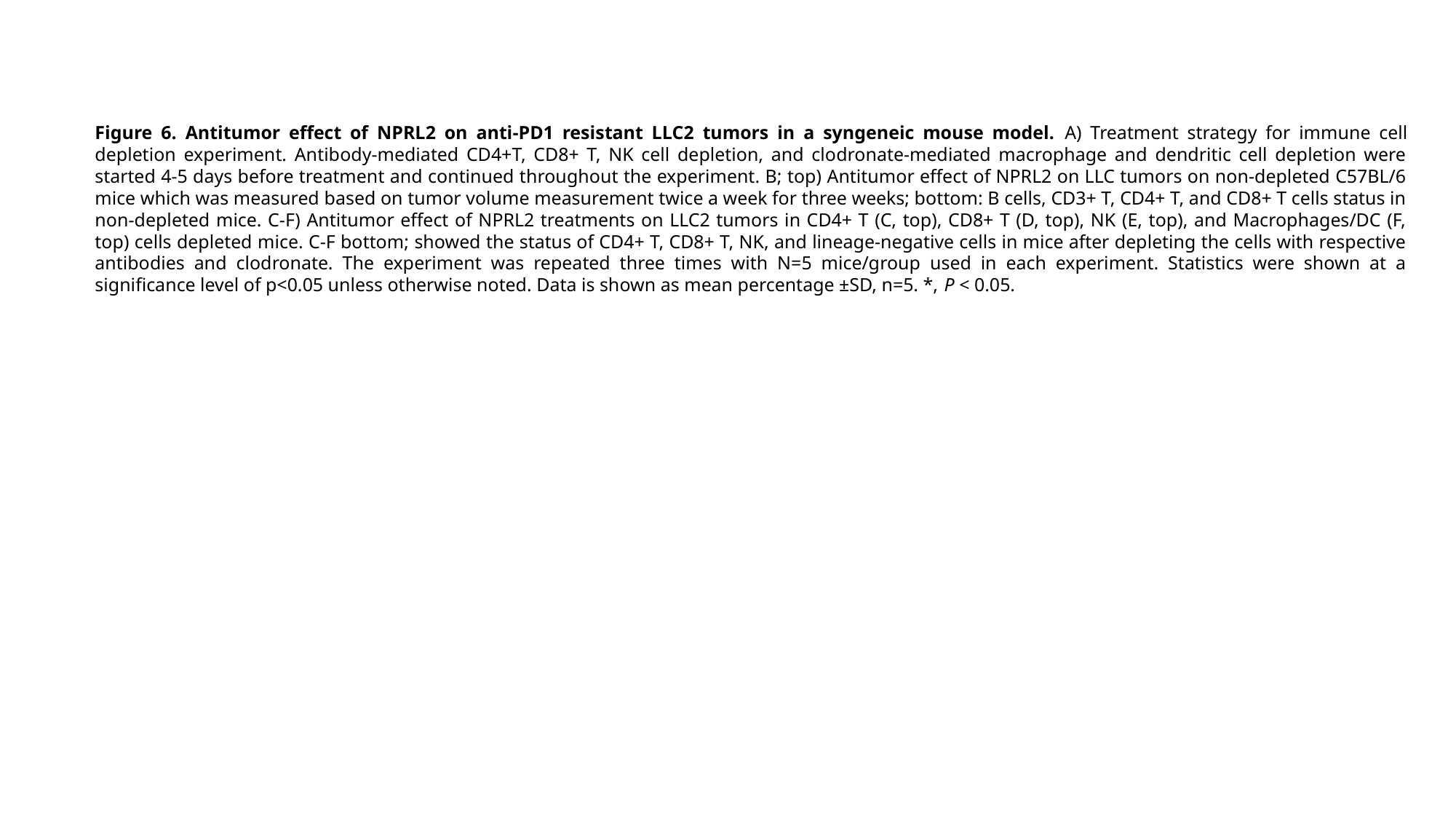

Figure 6. Antitumor effect of NPRL2 on anti-PD1 resistant LLC2 tumors in a syngeneic mouse model. A) Treatment strategy for immune cell depletion experiment. Antibody-mediated CD4+T, CD8+ T, NK cell depletion, and clodronate-mediated macrophage and dendritic cell depletion were started 4-5 days before treatment and continued throughout the experiment. B; top) Antitumor effect of NPRL2 on LLC tumors on non-depleted C57BL/6 mice which was measured based on tumor volume measurement twice a week for three weeks; bottom: B cells, CD3+ T, CD4+ T, and CD8+ T cells status in non-depleted mice. C-F) Antitumor effect of NPRL2 treatments on LLC2 tumors in CD4+ T (C, top), CD8+ T (D, top), NK (E, top), and Macrophages/DC (F, top) cells depleted mice. C-F bottom; showed the status of CD4+ T, CD8+ T, NK, and lineage-negative cells in mice after depleting the cells with respective antibodies and clodronate. The experiment was repeated three times with N=5 mice/group used in each experiment. Statistics were shown at a significance level of p<0.05 unless otherwise noted. Data is shown as mean percentage ±SD, n=5. *, P < 0.05.

### Slide 14
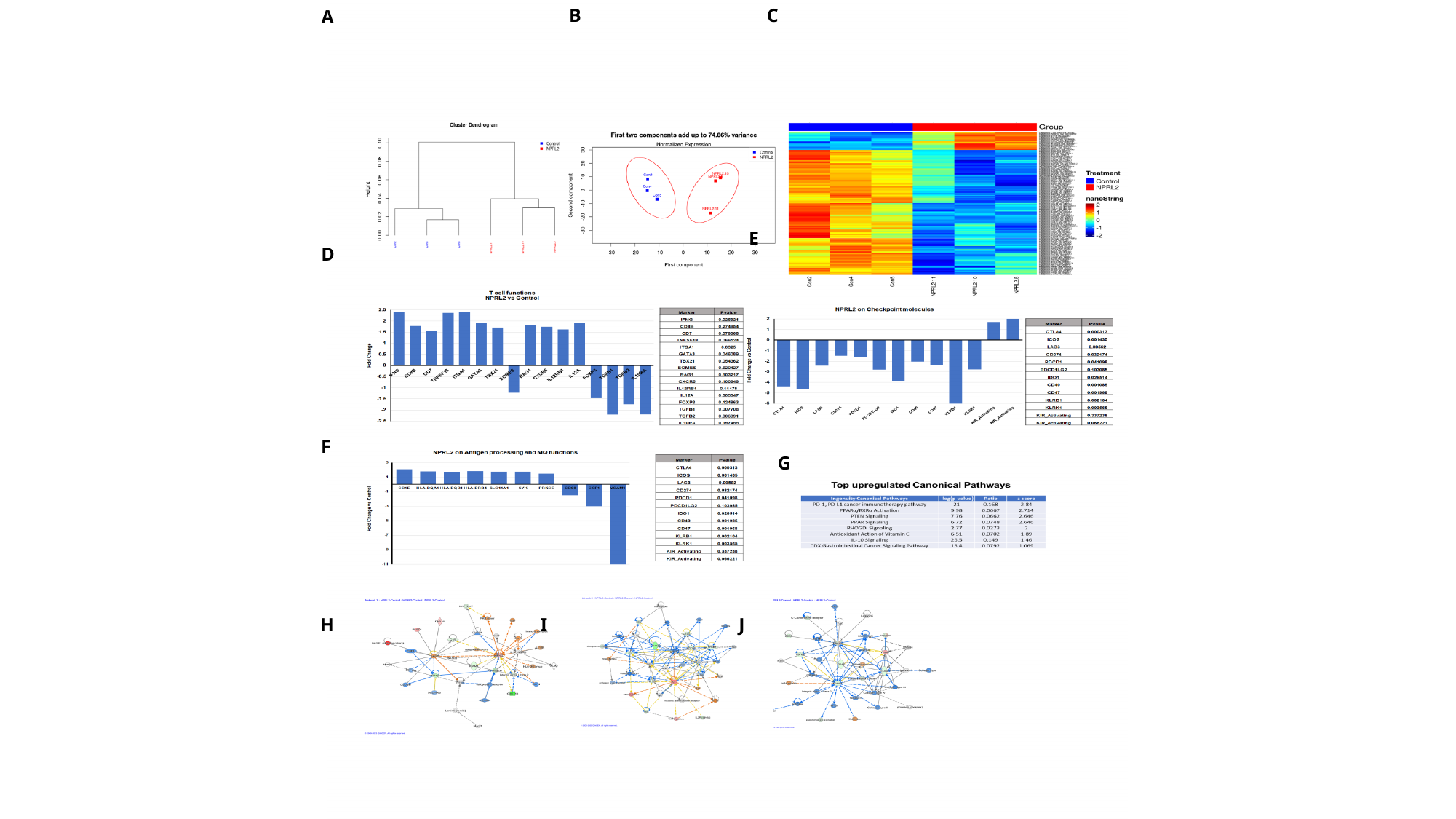

A
C
B
E
D
F
G
H
I
J

### Slide 15
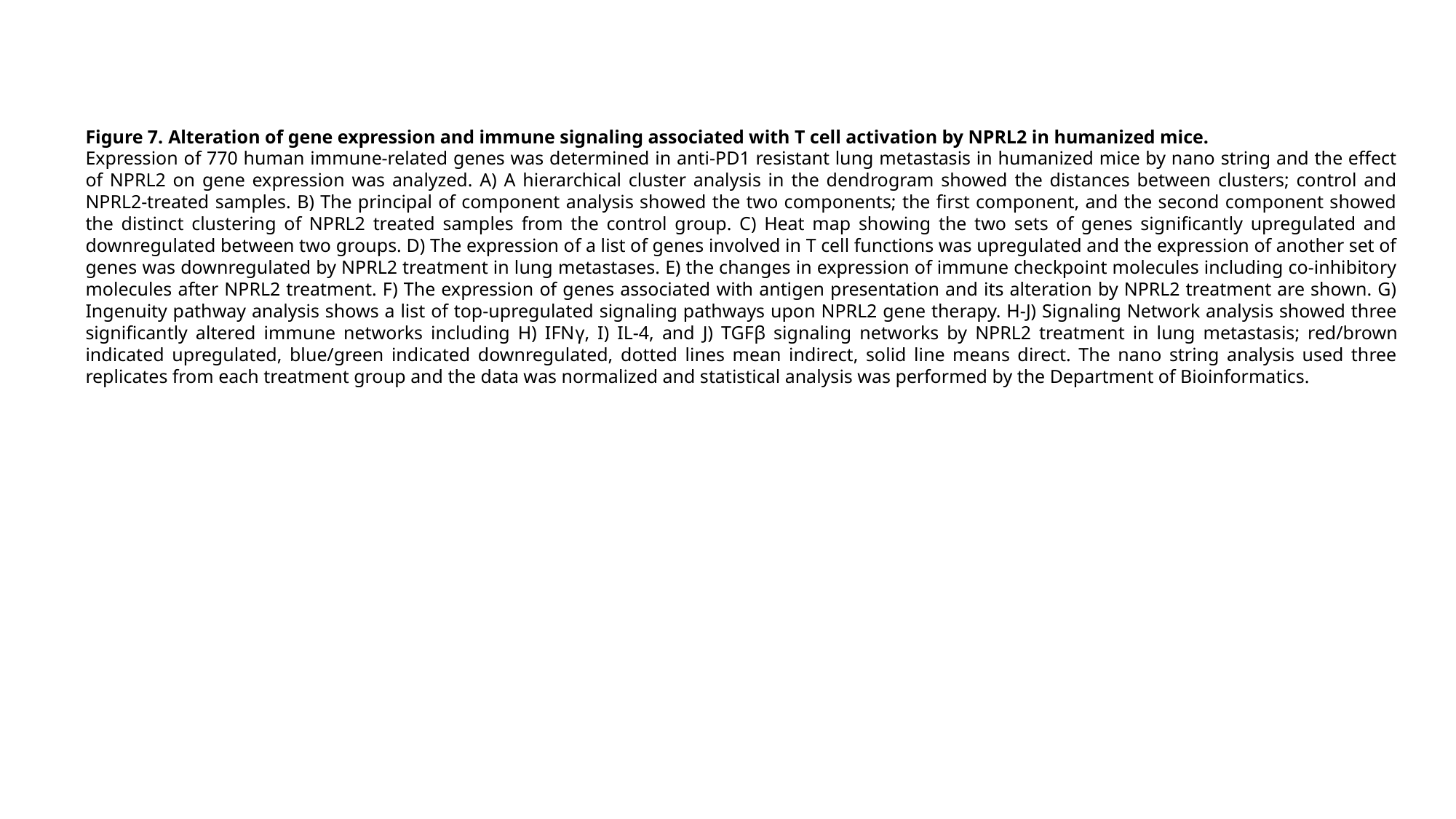

Figure 7. Alteration of gene expression and immune signaling associated with T cell activation by NPRL2 in humanized mice.
Expression of 770 human immune-related genes was determined in anti-PD1 resistant lung metastasis in humanized mice by nano string and the effect of NPRL2 on gene expression was analyzed. A) A hierarchical cluster analysis in the dendrogram showed the distances between clusters; control and NPRL2-treated samples. B) The principal of component analysis showed the two components; the first component, and the second component showed the distinct clustering of NPRL2 treated samples from the control group. C) Heat map showing the two sets of genes significantly upregulated and downregulated between two groups. D) The expression of a list of genes involved in T cell functions was upregulated and the expression of another set of genes was downregulated by NPRL2 treatment in lung metastases. E) the changes in expression of immune checkpoint molecules including co-inhibitory molecules after NPRL2 treatment. F) The expression of genes associated with antigen presentation and its alteration by NPRL2 treatment are shown. G) Ingenuity pathway analysis shows a list of top-upregulated signaling pathways upon NPRL2 gene therapy. H-J) Signaling Network analysis showed three significantly altered immune networks including H) IFNγ, I) IL-4, and J) TGFβ signaling networks by NPRL2 treatment in lung metastasis; red/brown indicated upregulated, blue/green indicated downregulated, dotted lines mean indirect, solid line means direct. The nano string analysis used three replicates from each treatment group and the data was normalized and statistical analysis was performed by the Department of Bioinformatics.

### Slide 16
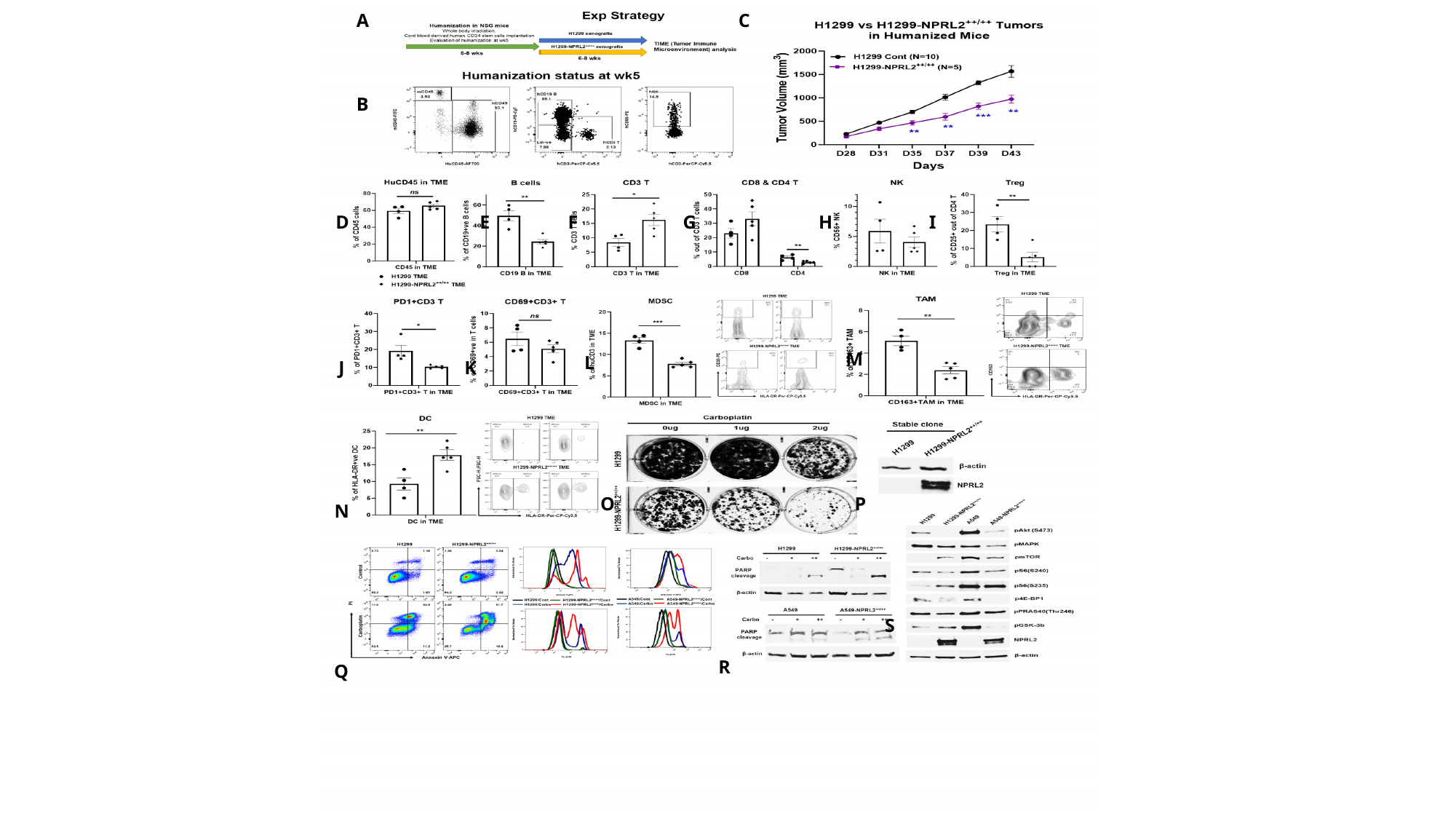

A
C
B
D
H
E
F
G
I
M
L
J
K
O
P
N
S
R
Q

### Slide 17
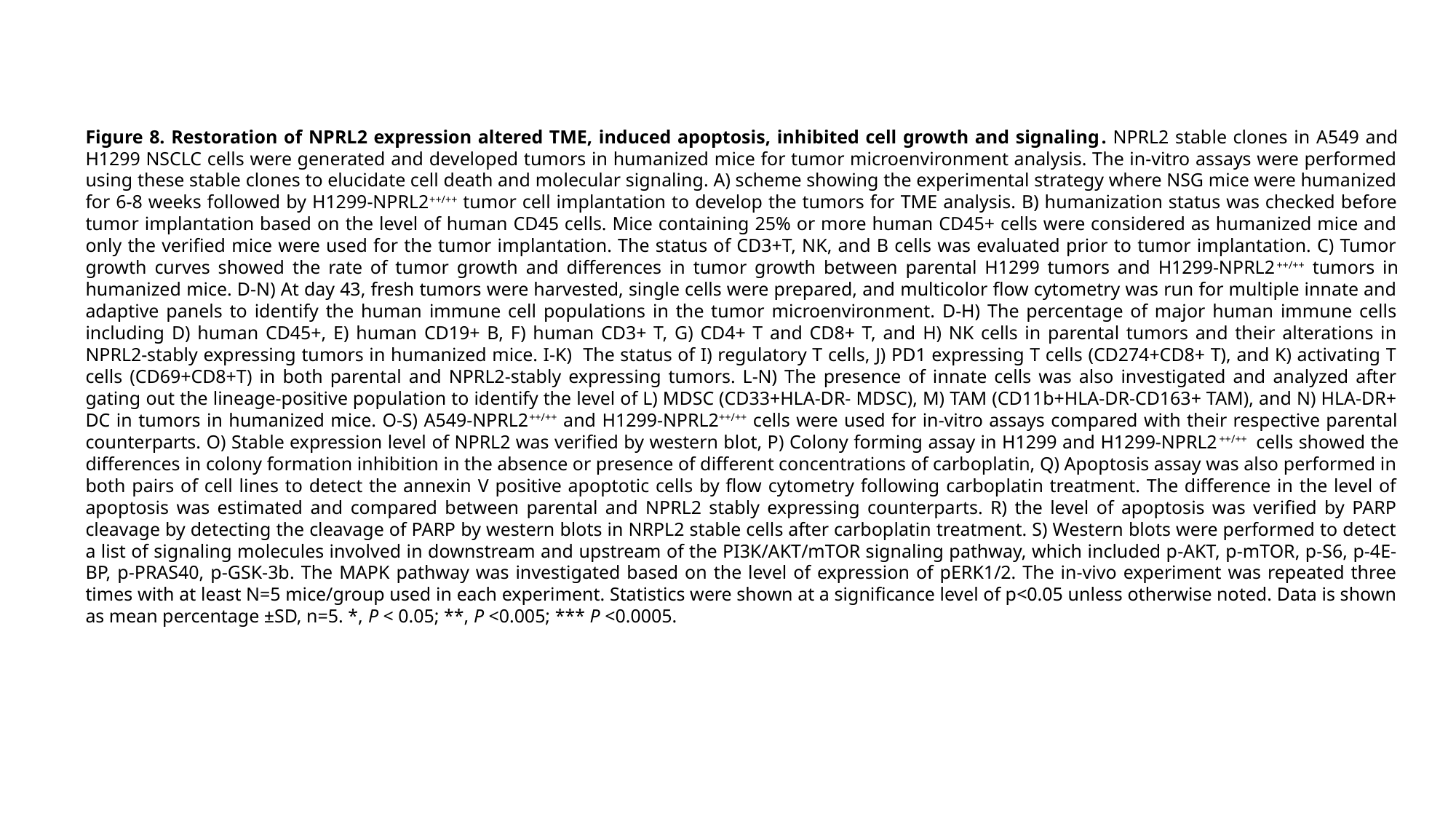

Figure 8. Restoration of NPRL2 expression altered TME, induced apoptosis, inhibited cell growth and signaling. NPRL2 stable clones in A549 and H1299 NSCLC cells were generated and developed tumors in humanized mice for tumor microenvironment analysis. The in-vitro assays were performed using these stable clones to elucidate cell death and molecular signaling. A) scheme showing the experimental strategy where NSG mice were humanized for 6-8 weeks followed by H1299-NPRL2^++/++^ tumor cell implantation to develop the tumors for TME analysis. B) humanization status was checked before tumor implantation based on the level of human CD45 cells. Mice containing 25% or more human CD45+ cells were considered as humanized mice and only the verified mice were used for the tumor implantation. The status of CD3+T, NK, and B cells was evaluated prior to tumor implantation. C) Tumor growth curves showed the rate of tumor growth and differences in tumor growth between parental H1299 tumors and H1299-NPRL2^++/++^ tumors in humanized mice. D-N) At day 43, fresh tumors were harvested, single cells were prepared, and multicolor flow cytometry was run for multiple innate and adaptive panels to identify the human immune cell populations in the tumor microenvironment. D-H) The percentage of major human immune cells including D) human CD45+, E) human CD19+ B, F) human CD3+ T, G) CD4+ T and CD8+ T, and H) NK cells in parental tumors and their alterations in NPRL2-stably expressing tumors in humanized mice. I-K) The status of I) regulatory T cells, J) PD1 expressing T cells (CD274+CD8+ T), and K) activating T cells (CD69+CD8+T) in both parental and NPRL2-stably expressing tumors. L-N) The presence of innate cells was also investigated and analyzed after gating out the lineage-positive population to identify the level of L) MDSC (CD33+HLA-DR- MDSC), M) TAM (CD11b+HLA-DR-CD163+ TAM), and N) HLA-DR+ DC in tumors in humanized mice. O-S) A549-NPRL2^++/++^ and H1299-NPRL2^++/++^ cells were used for in-vitro assays compared with their respective parental counterparts. O) Stable expression level of NPRL2 was verified by western blot, P) Colony forming assay in H1299 and H1299-NPRL2^++/++^ cells showed the differences in colony formation inhibition in the absence or presence of different concentrations of carboplatin, Q) Apoptosis assay was also performed in both pairs of cell lines to detect the annexin V positive apoptotic cells by flow cytometry following carboplatin treatment. The difference in the level of apoptosis was estimated and compared between parental and NPRL2 stably expressing counterparts. R) the level of apoptosis was verified by PARP cleavage by detecting the cleavage of PARP by western blots in NRPL2 stable cells after carboplatin treatment. S) Western blots were performed to detect a list of signaling molecules involved in downstream and upstream of the PI3K/AKT/mTOR signaling pathway, which included p-AKT, p-mTOR, p-S6, p-4E-BP, p-PRAS40, p-GSK-3b. The MAPK pathway was investigated based on the level of expression of pERK1/2. The in-vivo experiment was repeated three times with at least N=5 mice/group used in each experiment. Statistics were shown at a significance level of p<0.05 unless otherwise noted. Data is shown as mean percentage ±SD, n=5. *, P < 0.05; **, P <0.005; *** P <0.0005.

### Slide 18
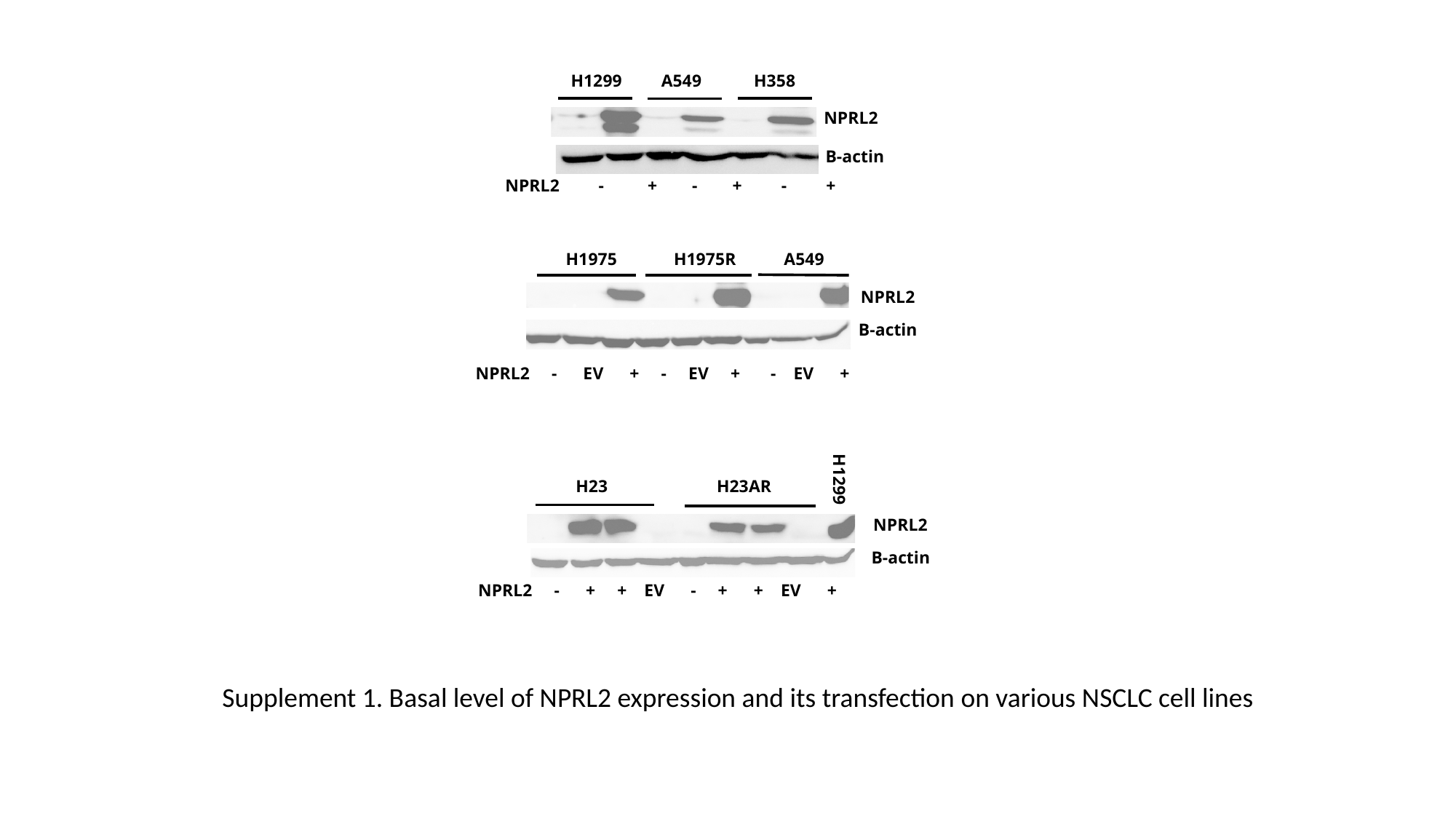

H1299 A549 H358
NPRL2
Β-actin
NPRL2 - + - + - +
H1975 H1975R A549
NPRL2
Β-actin
NPRL2 - EV + - EV + - EV +
H1299
H23 H23AR
NPRL2
Β-actin
NPRL2 - + + EV - + + EV +
Supplement 1. Basal level of NPRL2 expression and its transfection on various NSCLC cell lines

### Slide 19
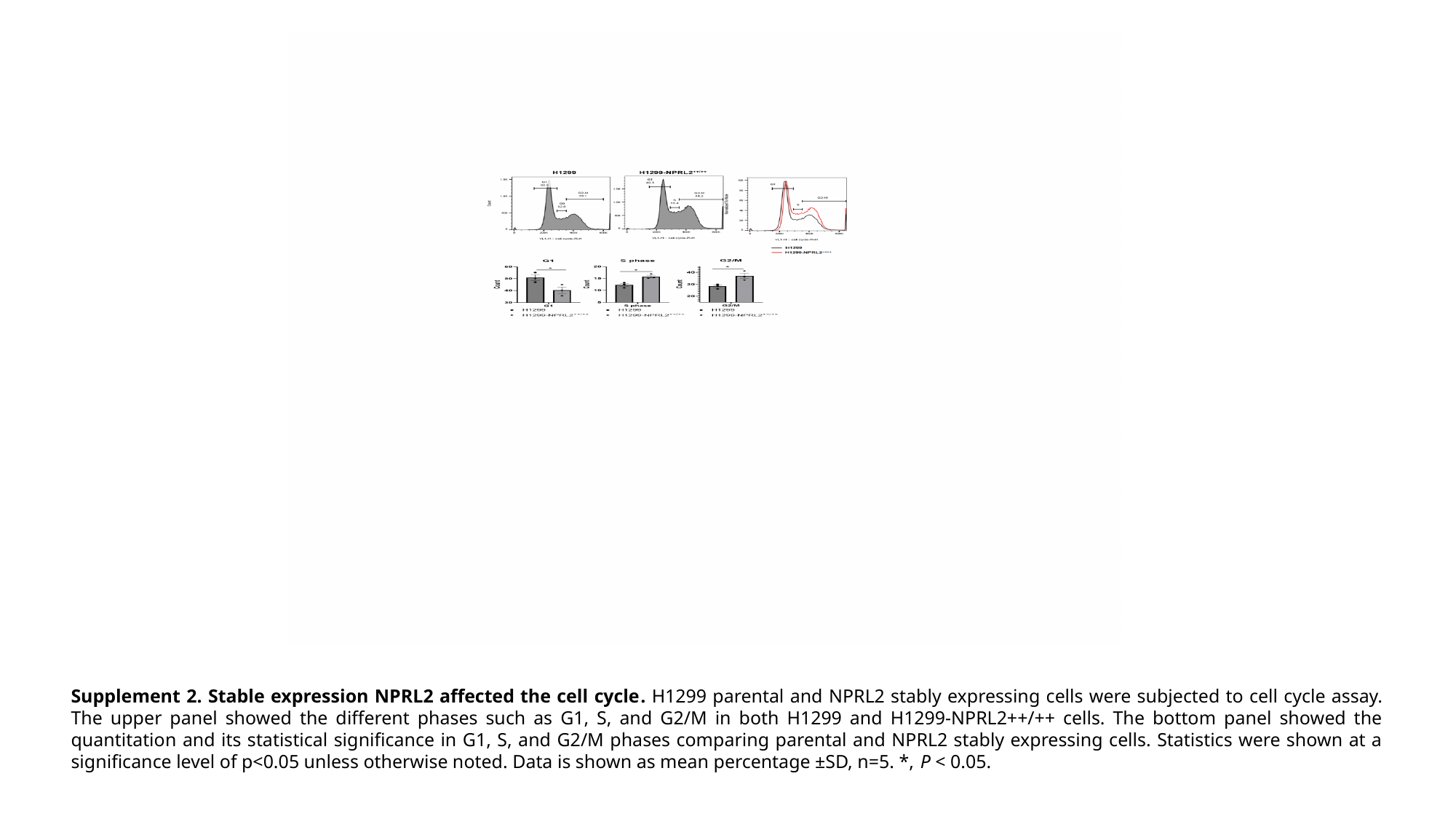

Supplement 2. Stable expression NPRL2 affected the cell cycle. H1299 parental and NPRL2 stably expressing cells were subjected to cell cycle assay. The upper panel showed the different phases such as G1, S, and G2/M in both H1299 and H1299-NPRL2++/++ cells. The bottom panel showed the quantitation and its statistical significance in G1, S, and G2/M phases comparing parental and NPRL2 stably expressing cells. Statistics were shown at a significance level of p<0.05 unless otherwise noted. Data is shown as mean percentage ±SD, n=5. *, P < 0.05.
